# Conservation, alteration, and redistribution of mammalian striatal interneurons

**DOI:** 10.1101/2024.07.29.605664

**Authors:** Emily K. Corrigan, Michael DeBerardine, Aunoy Poddar, Miguel Turrero García, Matthew T. Schmitz, Corey C. Harwell, Mercedes F. Paredes, Fenna M. Krienen, Alex A. Pollen

## Abstract

Mammalian brains vary in size, structure, and function, but the extent to which evolutionarily novel cell types contribute to this variation remains unresolved^1–4^. Recent studies suggest there is a primate-specific population of striatal inhibitory interneurons, the TAC3 interneurons^5^. However, there has not yet been a detailed analysis of the spatial and phylogenetic distribution of this population. Here, we profile single cell gene expression in the developing pig (an ungulate) and ferret (a carnivore), representing 94 million years divergence from primates, and assign newborn inhibitory neurons to initial classes first specified during development^6^. We find that the initial class of TAC3 interneurons represents an ancestral striatal population that is also deployed towards the cortex in pig and ferret. In adult mouse, we uncover a rare population expressing *Tac2*, the ortholog of *TAC3*, in ventromedial striatum, prompting a reexamination of developing mouse striatal interneuron initial classes by targeted enrichment of their precursors. We conclude that the TAC3 interneuron initial class is conserved across Boreoeutherian mammals, with the mouse population representing Th striatal interneurons, a subset of which expresses *Tac2*. This study suggests that initial classes of telencephalic inhibitory neurons are largely conserved and that during evolution, neuronal types in the mammalian brain change through redistribution and fate refinement, rather than by derivation of novel precursors early in development.

## Main

Ramón y Cajal first appreciated the abundance and morphological diversity of primate interneurons, linking these elaborations to “the functional superiority of the human brain”^7,8^. Cortical and striatal inhibitory neurons mainly emerge prenatally from progenitors lining the lateral ventricles of the ventral telencephalon, where spatial and temporal patterning signals influence the initial specification of inhibitory neuron subtypes^9,10^ and subsequent extrinsic cues further refine terminal class identity^11^. Striatal inhibitory neuron subtypes are generated primarily from embryonic progenitors expressing the transcription factor *NKX2.1* residing in the medial ganglionic eminence (MGE) and comprise terminal classes classically denoted by their expression of selective markers and neurochemical pathway genes, including *CHAT*-, *PTHLH, PVALB*-, *SST-,* and *TH*-expressing neurons^12–14^. In addition to these conserved classes, *TAC3*-expressing striatal interneurons constitute 30% of primate striatal interneurons, but were not observed in the first surveys of mouse and ferret striatum, raising the possibility that they may represent a derived primate-specific population^5^. Given the conservation of other initial classes of inhibitory neurons and regional transcription factor expression patterns during development^6,15^, we sought to examine the developmental specification of this class in additional species. Surprisingly, we find that initial classes of inhibitory neurons, including *TAC3,* are shared across Boreoeutherian mammals, with modifications of anatomical allocation and gene expression of derivatives from the TAC3 class in several taxa, suggesting that neural cell type evolution in mammals acts upon a conserved set of initial developmental classes.

### Conservation of initial classes in developing pig and ferret

We first assessed inhibitory neuron development and migration in the pig (*Sus scrofa*), a gyrencephalic mammal that diverged from the human lineage approximately 94 million years ago (**Fig. 1a**)^16^. Recent studies investigating porcine MGE development suggest that conserved transcription factors determine the fate and migration of interneurons^17^. We microdissected cortical and striatal regions from the pig brain at embryonic day 73 (E73), a stage comparable to the second trimester of human development (**Fig. 1b)**, to capture cells during peak interneuron specification^18–20^, and performed single cell RNA sequencing (scRNAseq) using 10x Genomics Next GEM Single Cell 3’ HT v3.1 technology. We applied stringent quality control metrics, performed dimensionality reduction, batch correction, and Leiden clustering on 25,089 cells. To focus our investigation of inhibitory neurons, we isolated this population using the conserved markers *DLX*1/2/5/6 and *GAD1/2* (methods), and performed Leiden clustering on 8,140 inhibitory cells (**Extended Data Fig. 1a**,b). We next classified these clusters based on region-specific transcription factors indicating MGE (*NKX2.1*, *LHX6*), LGE (*PAX6*, *MEIS2*, *FOXP2*), or CGE (*NR2F2*)^9,21^ origin (**Extended Data Fig. 1c-g**), and markers of initial classes defined by recent macaque and mouse data^6^. Of 11 initial classes previously observed in the developing macaque, we recovered 9 in developing pig based on expression of region-specific transcription factors and shared markers (**Fig. 1c, d**). We did not observe a distinct MGE-derived LHX6/NPY-expressing cluster indicative of the MGE_LHX6/NPY initial class found in primates and rodents, which was likely due to undersampling in pig (**Extended Data Fig. 1c)**.

**Figure 1:**
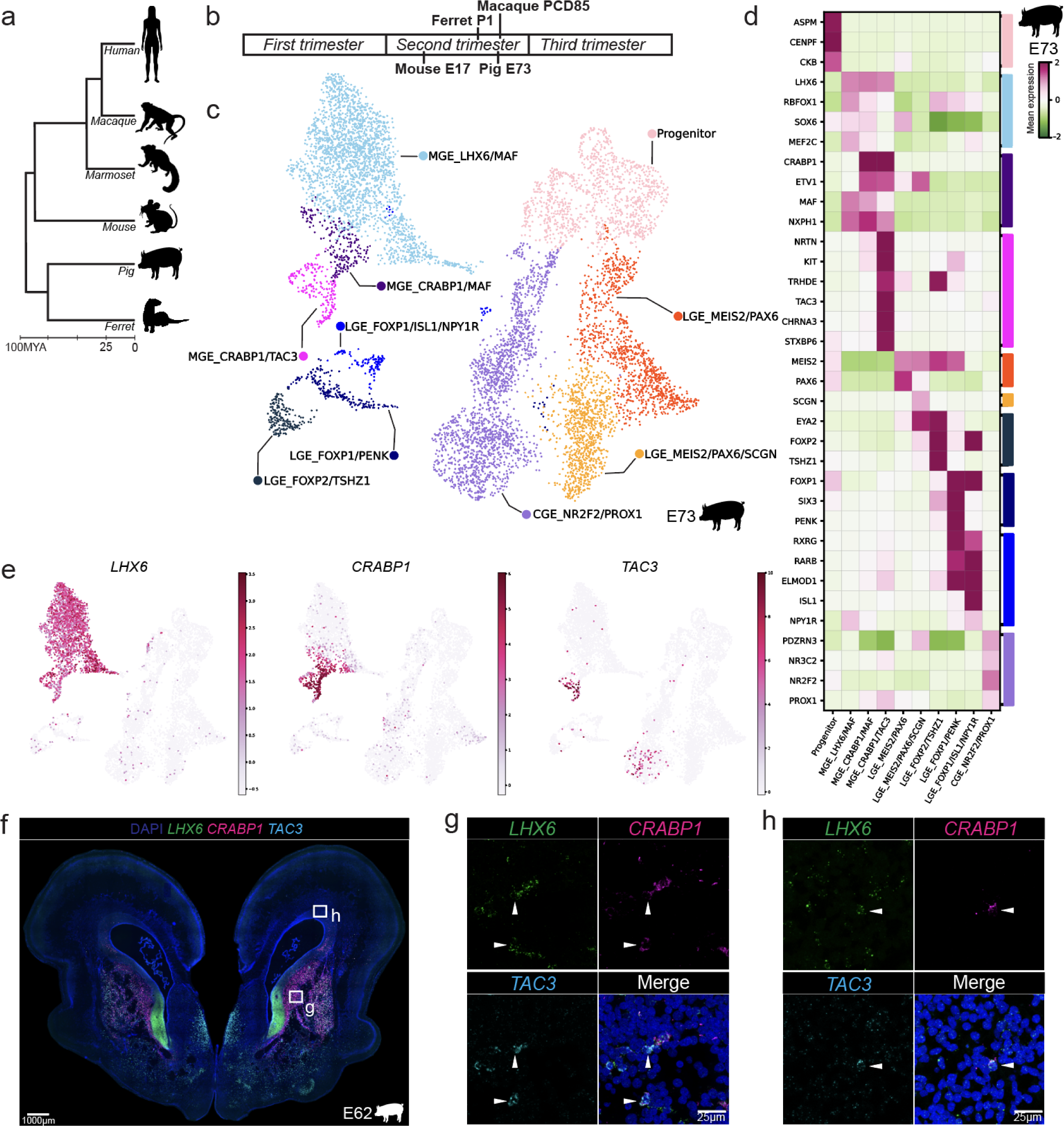
Survey of inhibitory neuron initial classes reveals the MGE_CRABP1/TAC3 class is conserved and redistributed in the developing pig brain. **a.** Taxonomy of selected Boreoeutherian species. **b.** Timing of species surveyed during development compared to human development. **c.** UMAP colored by inhibitory neuron initial classes in the E73 pig. **d.** Heatmap of selected markers of inhibitory neuron initial classes in developing pig, scaled and normalized. **e.** UMAP colored by expression of MGE_CRABP1/TAC3 initial class marker genes in pig, scaled and normalized. **f.** Whole section image acquired at 10X of E63 developing pig RNAscope with markers *LHX6*, *CRABP1* and *TAC3*, with boxes highlighting the MGE_CRABP1/TAC3 class in striatum (g), and cortex (h). **g,h.** High magnification (100X) maximum intensity projection with arrowheads indicating the MGE_CRABP1/TAC3 initial class.

Previous studies suggest that the conserved MGE_CRABP1/MAF population gives rise to PVALB, PTHLH, and TH striatal interneurons in mice and primates^13^, while the related initial class, MGE_CRABP1/TAC3, gives rise to TAC3 neurons in primates^6^. In pig, we observed the conserved MGE_CRABP1/MAF class, but unexpectedly, we also identified a distinct cluster (191 cells) that expresses markers of the primate MGE_CRABP1/TAC3 class such as *NKX2.1*, *LHX6*, *CRABP1*, *CHRNA3*, and *TAC3* (**Fig. 1d,e**). To confirm the spatial location of this initial class in developing pigs, we performed RNA *in situ* hybridization (RNAscope) at E62 (**Fig. 1f**). We observed a *LHX6*, *CRABP1*, *TAC3* positive population in the striatum (**Fig. 1g**) of developing pig, revealing that the MGE_CRABP1/TAC3 initial class is conserved in developing pig striatum.

Analysis in marmoset, human, and macaque suggests that cells derived from the MGE_CRABP1/TAC3 initial class are present in the primate striatum but not neocortex^5,6,22^. Unexpectedly, we observed a novel population of *LHX6*, *CRABP1*, *TAC3* expressing cells migrating towards (**Extended Data Fig. 2a,b**) and within (**Fig. 1h**) the cortex of E62 pig. We performed RNAscope on E73 developing pig and again observed this MGE_CRABP1/TAC3 initial class in the striatum and cortex (**Extended Data Fig. 2c-f**). Finally, we performed RNAscope on the adult pig and confirmed that these populations persist past development (**Extended Data Fig. 2g-i**), indicating the MGE_CRABP1/TAC3 initial class is also deployed to the cortex in the developing pig.

To determine if the MGE_CRABP1/TAC3 class is present in other Boreoeutherian clades, we next evaluated the presence of TAC3 neurons in the ferret, a carnivore that diverged from primates approximately 94 M years ago and from pig approximately 76 M years ago (**Fig. 1a**). Although early surveys in adult ferret did not detect TAC3 neurons among other striatal populations^5^, we further examined the developing ferret brain, given the presence of this initial class in the pig and the conserved features of cortical development between ferret and primate^23^. We performed scRNAseq on the cortex and subpallium of postnatal day 1 (P1) developing ferret (*Mustella putorius furo*), a comparable stage of inhibitory neuron production to E62 in pig. We again applied stringent quality control metrics and performed Leiden clustering on 21,493 cells. We then isolated 10,540 cells from the inhibitory lineage, renormalized and reclustered (**Extended Data Fig. 3a,b**), and classified cell types based on the expression of conserved marker genes in mouse and primate, revealing 12 conserved initial classes (**Fig. 2a-c, Extended Data Fig. 3c-g**). We did not recover a distinct MGE_CRABP1/MAF initial class. However, the inability to detect MGE_CRABP1/MAF is likely due to low cell coverage, since the *PTHLH* and *PVALB*-expressing derivatives of this class are present in adult ferret^5^. As in pig, we again observed a cluster 161 cells) expressing the gene signatures of MGE_CRABP1/TAC3 (**Fig. 2b,c**), suggesting that TAC3 neurons are derived from an initial class already present in the common ancestor of Boroeutherians.

**Figure 2:**
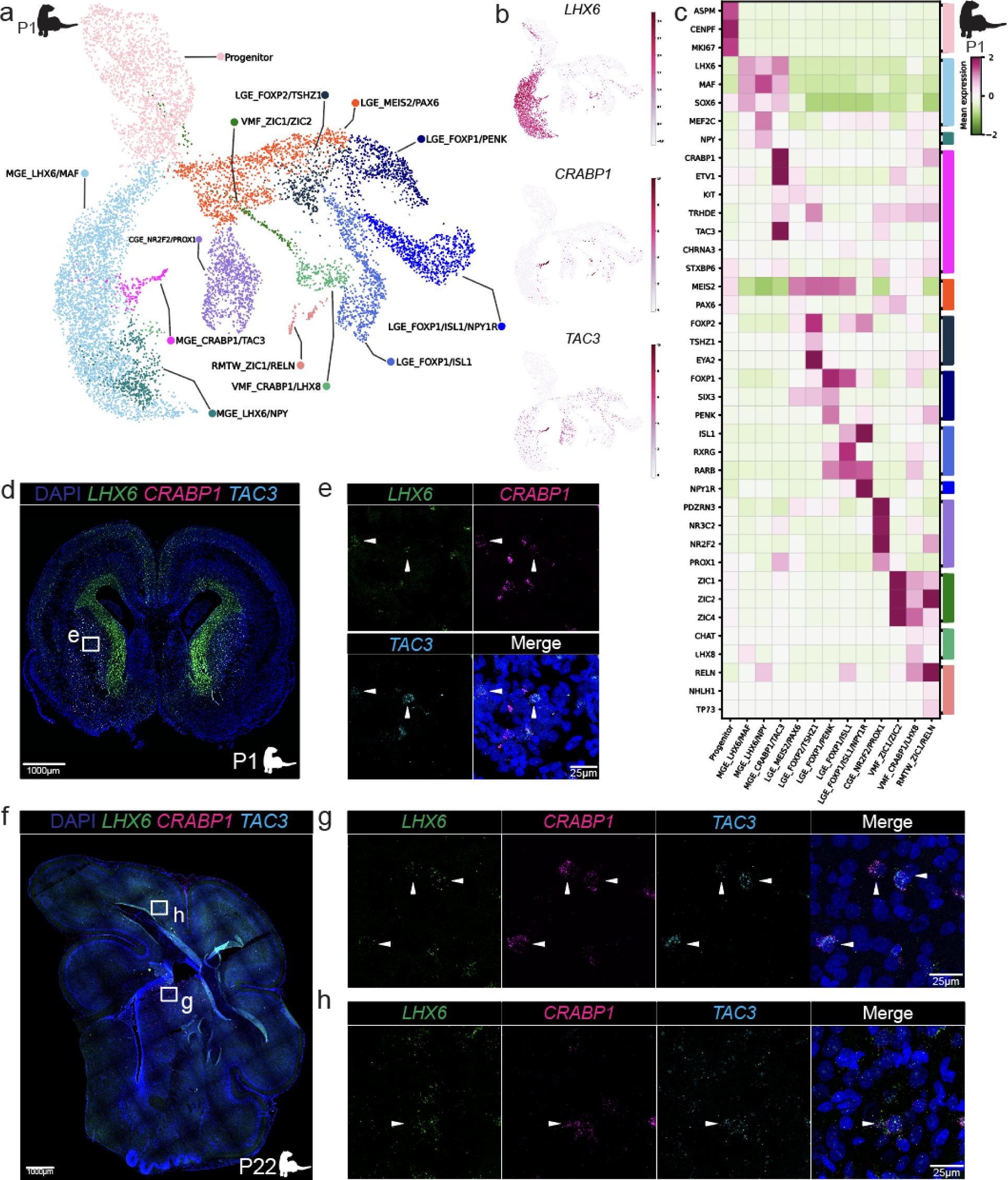
The MGE_CRABP1/TAC3 class is conserved and redistributed in Laurasiatherians. **a.** UMAP colored by inhibitory neuron initial classes in the P1 developing ferret brain. **b.** UMAP colored by expression of MGE_CRABP1/TAC3 initial class marker genes, scaled and normalized. **c.** Heatmap of selected markers of inhibitory neuron initial classes in developing ferret, scaled and normalized. **d.** Whole section image acquired at 20x of P1 developing ferret RNAscope with markers *LHX6*, *CRABP1*, and *TAC3,* and box highlighting the MGE_CRABP1/TAC3 class in striatum (e). **e.** High magnification (100X) maximum intensity projection with arrowheads indicating the MGE_CRABP1/TAC3 class. **f.** Whole section image acquired at 10X of RNAscope in the P22 ferret brain, with the markers *LHX6*, *CRABP1*, and *TAC3*, and boxes highlighting striatum (g) and cortex (h). **g, h.** High magnification (100X) maximum intensity projection with arrowheads indicating MGE_CRABP1/TAC3 class.

To examine the distribution of the MGE_CRABP1/TAC3 initial class in developing ferrets, we performed RNAscope with the markers *LHX6*, *CRABP1*, and *TAC3*. We found a *LHX6, CRABP1, TAC3* positive population in the striatum of the P1 developing ferret (**Fig. 2d,e**). We did not observe MGE_CRABP1/TAC3 in or migrating towards the cortex of the ferret at P1. However, we performed RNAscope on P22 ferret and observed the striatal MGE_CRABP1/TAC3 population as well as MGE_CRABP1/TAC3 cells in the cortex, suggesting that the cortically-bound population may only migrate after P1(**Fig. 2f-h**). Finally, we confirmed the presence of these populations in the adult cortex and striatum (**Extended Data Fig. 4a-c**)

Given the presence of the MGE_CRABP1/TAC3 initial class in the cortex of both pig and ferret, we further examined the distribution of this initial class in developing and adolescent macaque. Recent studies described CRABP1 cells migrating to the cortex in the developing human and linked these to a cortical PVALB population^15^. We performed RNAscope in post-conception day (PCD) 65, PCD80 and 7 month old macaque and did not observe any cells coexpressing *LHX6*, *CRABP1*, and *TAC3* in the cortex (**Extended Data Fig. 4d-i**). Similarly, we analyzed gene expression of 4,989 *LHX6 CRABP1* expressing cells from PCD37-110 developing macaque across the telencephalon^24^ and only detected MGE_CRABP1/TAC3 neurons in dissections that include striatum (**Extended Data Fig. 5a-f**). These results indicate that MGE_CRABP1/TAC3 interneurons are conserved in the striatum of primate, ferret, and pig, and migrate to the cortex in pig and ferret.

### Comparative analysis of terminal classes

Our observation of a conserved MGE_CRABP1/TAC3 initial class in ferret, pig (this study) and primate raises the question of whether this class is truly lost in mice. Our previous studies examining both adult and developing mouse data did not detect TAC3 interneurons, nor a MGE_CRABP1/TAC3 initial class after examining over 140,000 adult mouse inhibitory neurons and 76,000 developing mouse inhibitory neurons^6^. However, a recently published atlas of 4 million sequenced cells provides an order of magnitude greater resolution into the cell composition of the adult mouse brain^25^. Using this newly published dataset, we isolated 3,586 striatal inhibitory interneurons, and reclustered and classified the populations based on known markers (**Fig. 3a, Extended Data Fig. 6a,b**) (see Methods).

**Figure 3:**
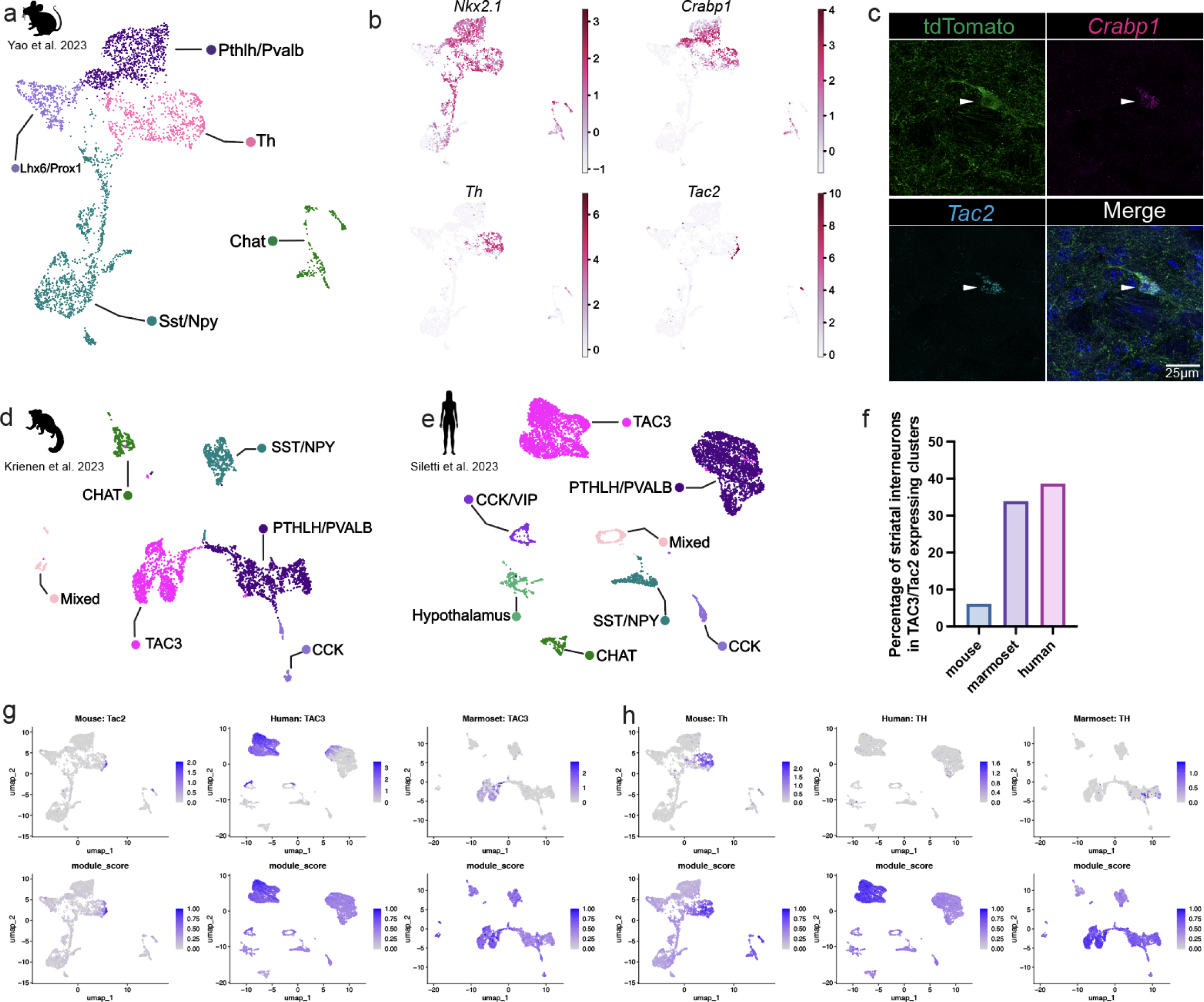
Identification of rare *Tac2*-expressing population in adult mouse TH interneurons. **a.** UMAP colored by striatal inhibitory neuron terminal classes in adult mouse. **b.** UMAP colored by markers of Th interneuron class, scaled and normalized. **c.** High magnification (100X) maximum intensity projection of adult Nkx2.1-Cre;Ai14 mouse brain tdTomato reporter expression with *Crabp1* and *Tac2* RNAscope and arrowheads indicating a rare Tac2 interneuron in the mouse. **d.** UMAP colored by striatal inhibitory neuron terminal classes in adult marmoset. **e.** UMAP colored by striatal inhibitory neuron terminal classes in adult human. **f.** Percentage of cells in Tac2/TAC3 clusters out of total striatal inhibitory neurons (mouse: n = 3,586, marmoset: n= 3,281), human: n= 6,042). In mouse, Allen institute parent clusters 0841 and 0842 are plotted which contain a subset of Tac2-expressing cells. **g.** Top row: UMAP colored by *Tac2*/*TAC3* expression in mouse, human, and marmoset datasets. Bottom row: Hotspot mouse *Tac2* gene module score when projected onto mouse, human and marmoset data. **h.** Top row: UMAP colored by *Th* expression in mouse, human, and marmoset datasets. Bottom row: Hotspot mouse *Th* gene module scores when projected onto mouse, human and marmoset data.

In mouse, striatal Th interneurons are a morphologically and physiologically diverse class that connects both to principal spiny projection neurons and to a subset of other striatal interneuron classes^26^. Despite expressing tyrosine hydroxlyase (*Th*), they do not appear to synthesize dopamine^26,27^. Within the mouse *Th*-expressing cluster (676 cells), we observed a small population which expressed characteristic markers of TAC3 interneurons such as *Crabp1*, *Tac2*, *Chrna3*, and *Stxbp6* (**Fig. 3b**) and confirmed the presence of *Nkx2.1, Crabp1, Tac2* co-expressing cells by RNAscope (**Fig. 3c**). Mouse *Tac2*-expressing cells were mainly found within the STR Lhx8 Gaba_1 clusters 0841 and 0842 as annotated by the Allen Brain Cell Atlas (**Extended Data Fig. 6c,d)**. These parent clusters are localized to the ventromedial striatum and septum, mirroring the sparse ventromedial expression of *Tac2* within Th-striatal interneurons (**Extended Data Fig. 6e**).

In contrast to primates, where TAC3 interneurons represent ∼30% of striatal interneurons, the parent clusters (*Th*+) containing *Tac2* expressing cells accounted for 6.1% of mouse striatal interneurons (of which 51 cells, or 1.4% of striatal interneurons, expressed *Tac2*) (**Fig. 3d-f**). Analysis of BAX null mice with impaired apoptosis revealed that cell death could not account for the reduced fraction of the *Tac2*-expressing population in mouse (**Extended Data Fig. 6f**).

To investigate cell type homology between mouse *Tac2*-expressing Th neurons and primate TAC3 neurons, we isolated striatal interneurons from published adult human^28^ and marmoset^22^ datasets (**Fig. 3d,e**) (see Methods). We observed major striatal interneuron types SST/NPY, CHAT, PTHLH/PVALB, CCK, and TAC3 within both primate datasets. In human, we also recovered CCK/VIP interneurons, a *TAC3*-expressing presumptive CGE-derived population (based on *ADARB2* expression) that is distinct from MGE-derived TAC3 neurons. Within primate striatal interneurons, *TH* was mainly detected in a subset of the PTHLH/PVALB population (**Extended Data Fig. 7a-d**), a derivative of the CRABP1/MAF class, and exhibited limited expression in TAC3 interneurons in contrast to mouse, raising the possibility that *TH* marks different subtypes in rodents and primates.

Next, we co-embedded the adult human^28^, marmoset^22^, and mouse^25^ striatal interneurons in a shared latent space using LIGER^29^. We evaluated cross-species relationships by the frequency of overlap between cell types from each species in integrated clusters across 126 iterations covering a wide range of parameter combinations related to feature space, dimensionality, and the tolerance for species-specific clusters (see Methods). For comparisons of marmoset and human, PTHLH/PVALB, SST/NPY, TAC3, CHAT, and CCK populations were most frequently observed co-embedding with their expected counterpart. Comparisons of mouse to both marmoset and human supported homology between the rare Tac2 population and primate TAC3 interneurons, with some evidence for correspondence of the broader mouse Th+ Tac2-populations to primate TAC3 neurons as well (**Extended Data Fig. 7e-g**). We also observed variability in co-embedding at this phylogenetic distance, including mouse Chat cells mapping to both marmoset and human TAC3 and CHAT and limited correspondence between mouse and human SST/NPY populations (**Extended Data Fig. 7e-l**).

To further examine the relationship between the terminal classes across species, we projected gene coexpression modules that mark interneuron populations in one species onto other species using Hotspot^30^. In each species, Hotspot was used to partition genes into non-overlapping modules, and we subsequently identified the modules most correlated with individual marker genes of interest (**Supplementary Table 1**). Gene modules marking *Sst*- and *Chat-* populations in mouse showed strong enrichment in the corresponding primate cell types, as did reciprocal projection of primate modules to mouse (**Extended Data Fig. 8a-f**). Projecting the mouse *Tac2* module onto both human and marmoset datasets further supported homology with primate TAC3 interneuron populations (**Fig. 3g**). In addition, the mouse *Th* module also projected to the primate TAC3 interneurons (**Fig. 3h**), consistent with the recent observation of shared marker genes in mouse Th and human TAC3 populations^31^. In contrast, the marmoset *TH* module mapped to human and mouse PTHLH/PVALB, supporting a distinction between primate and rodent *TH* populations(**Extended Data Fig. 8g**). However, reciprocal projections of human and marmoset *TAC3* modules showed strongest correspondence to the mouse Sst/Npy class with some expression in the mouse *Tac2* and Th populations (**Extended Data Fig. 8h-j**). Together, analysis of terminal classes in adult scRNA-seq data revealed a rare *Tac2*-expressing population enriched in mouse ventromedial forebrain with evidence for homology to primate TAC3 counterparts, and distinctions between mouse and primate TH interneurons, but the homologies were difficult to determine by analysis of adult data alone.

### Modification of conserved class in mouse

To resolve the possible homologies among terminal classes, we considered three scenarios during development that could lead to the rare *Tac2*-expressing population in the adult mouse and differences in TH expression between primate and rodents (**Extended Data Fig. 9a**). First, the MGE_CRABP1/TAC3 class could have been lost in mouse, and the small Tac2 mouse population and Th-interneurons could be independently produced from the MGE_CRABP1/MAF class. Second, the MGE_CRABP1/TAC3 initial class could exist in mouse but as a small population, giving rise to the rare Tac2 population in adult mouse, with the mouse MGE_CRABP1/MAF class producing Th cells. Finally, there could be two *Crabp1* initial classes of comparable abundance in mouse, one of which gives rise to the Pthlh and Pvalb populations, while the other gives rise to the Th and small Tac2 populations, consistent with the observed transcriptional similarity of these populations in adult mouse. To distinguish between these hypotheses, we performed high cellular coverage analysis of embryonic mouse striatal interneuron development by enriching for MGE-derived populations.

To enrich for striatal initial classes in developing mouse, we dissected the striatum from Nkx2.1-cre/Ai14 mouse embryos at E15, E17 and E18, sorted for cells with a developmental history of *Nkx2.1* expression (indicative of MGE origin), and performed single cell sequencing on that population (**Fig. 4a**). We isolated inhibitory neurons and their progenitors, performed dimensionality reduction, batch correction, and Leiden clustering on 71,036 cells, and classified these clusters based on known makers (**Fig. 4b-d, Extended Data Fig. 9**). We recovered predominantly MGE-derived inhibitory neurons, marked by *Lhx6* and *Nkx2.1*, including two distinct *Crabp1* populations of comparable abundance, each containing cells from all three developmental stages surveyed (**Extended Data Fig. 9c**). One Crabp1 population (2,353 cells) coexpressed the markers *Maf* and *Rbp4*, indicating it represented the conserved MGE_CRABP1/MAF initial class. The second Crabp1 positive population (1,393 cells) expressed the markers *Chrna3*, *Stxbp6, Trh* and high levels of *Th*, with a small subset (7 cells) also expressing *Tac2* (**Fig. 4c**). This observation of *Tac2* gene expression in a fraction of cells in the Crabp1/Th cluster resembled the pattern observed in adult mouse data. In contrast, *TH* was not detected in the CRABP1/TAC3 population in primates and Laurasiatherians (**Extended Data Fig. 10**). We performed RNAscope on developing (E17) and adult mouse tissue using the markers *Nkx2.1*, *Crabp1*, and *Chrna3*, and observed this population only in the striatum (**Fig. 4e, Extended Data Fig. 10a**).

**Figure 4:**
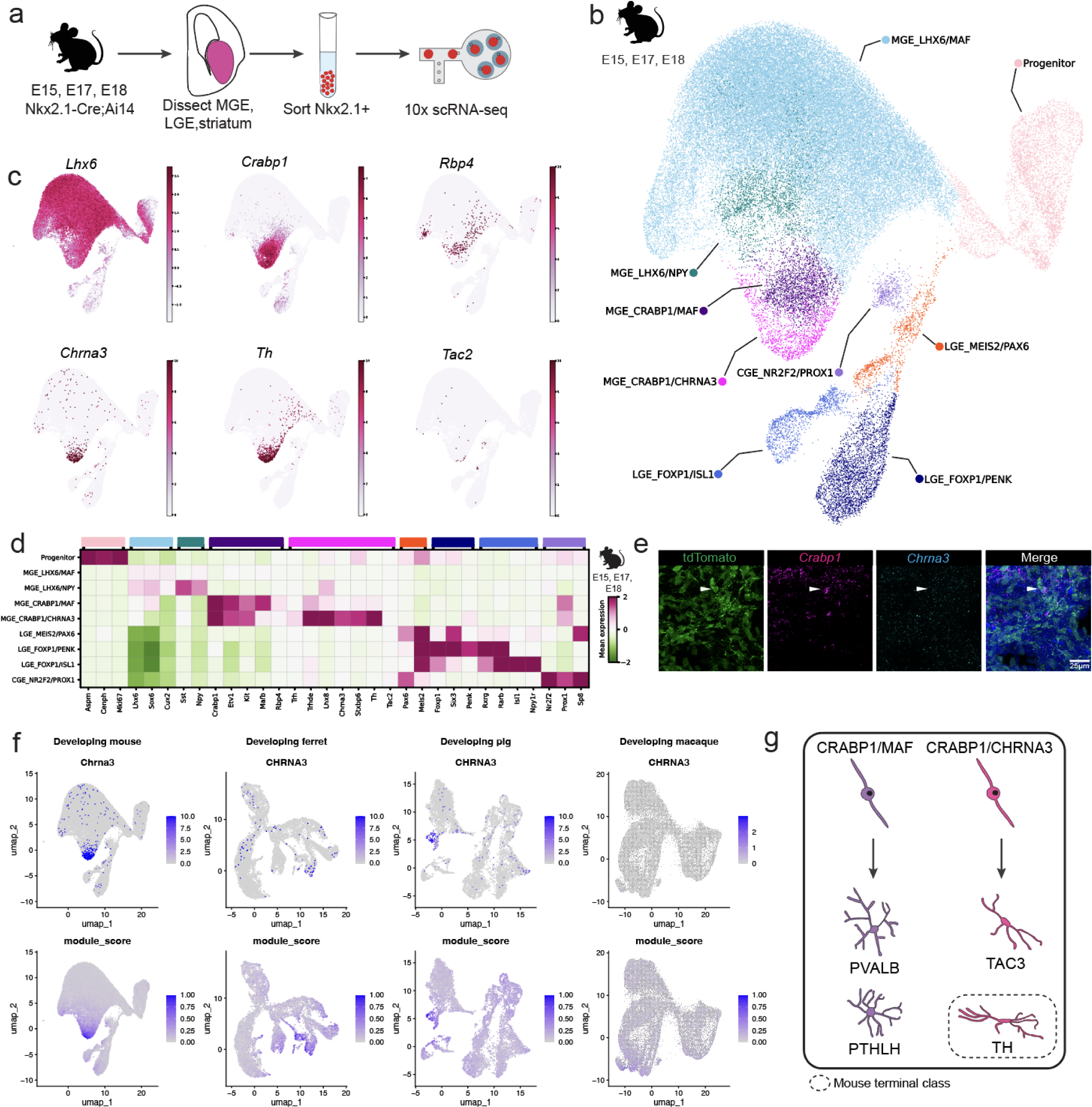
TAC3 initial class has been altered in mouse to produce Th striatal interneurons. **a.** Schematic of experimental design. **b.** UMAP colored by inhibitory neuron initial classes recovered from developing mouse MGE, LGE and striatum. **c.** UMAP colored by markers of *Lhx6*/*Crabp1* initial classes, scaled and normalized. **d.** Heatmap of selected markers of recovered inhibitory initial class markers, scaled and normalized. **e.** High magnification (100X) maximum intensity projection of E17 Nkx2.1-Cre;Ai14 developing mouse brain tdTomato reporter expression and *Crabp1*, *Chrna3* RNAscope with arrowhead indicating the MGE_CRABP1/CHRNA3 initial class. **f.** Top row: UMAP colored by *CHRNA3* expression in mouse, ferret, pig and macaque datasets inhibitory interneuron datasets. Bottom row: Hotspot mouse *Chrna3* gene module projected onto developing mouse, ferret, pig, and macaque datasets. **g.** Schematic summarizing *LHX6*/*CRABP1* initial and terminal classes in Boreoeutherian mammal striatum.

Finally, we utilized Hotspot to examine homology between the *Th* or *TAC3 CRABP1-*expressing initial classes in developing mouse, pig, ferret and published macaque^6^ (**Extended Data** Fig 10c,d) inhibitory interneuron datasets. Mouse markers including *Chrna3, Tac2,* and *Th* comprised one module which marked the *Crabp1/Th* cluster in the mouse and corresponded to the MGE_CRABP1/TAC3 population in macaque and pig, with the highest module score in the VMF_CRABP1/LHX8 population in ferret (**Fig. 4f, Extended Data Fig. 10**). Reciprocally, pig markers including *TAC3* and *CHRNA3* comprised a module that marked the pig MGE_CRABP1/TAC3 cluster and projected to ferret and macaque MGE_CRABP1/TAC3 clusters, and mouse *Crabp1*/*Th* population (**Extended Data Fig. 10, Supplementary Table 1)**. These findings indicate that both initial classes of CRABP1 striatal interneurons observed in primates, pig, and ferret are conserved in mouse, with the mouse population showing a derived, but not complete, loss of *Tac2* expression, and a gain of *Th* expression early in neuronal development that persists in adulthood (**Extended Data Fig. 10b**).

Based on the expression of the conserved markers *Crabp1* and *Chrna3* in the *Th* (mouse) or *TAC3*-expressing CRABP1 initial class across most species sampled, we conclude that there are two conserved initial classes of *LHX6*/*CRABP1*-expressing striatal interneuron initial classes across Boreoeutherian mammals, the MGE_CRABP1/MAF and the MGE_CRABP1/CHRNA3 class, which modifies its neuropeptide or neuromodulator gene expression as well as destination throughout evolution (**Fig. 4g, Extended Data Fig. 10k**).

## Discussion

By exploring developmental and adult cell type diversity across a wider range of taxa, we find that mouse Th striatal interneurons, previously considered to be homologous to primate TH striatal interneurons^27^, instead represent a modified derivative of TAC3 interneurons, with both arising from an ancestral initial class in Boroeutherians. We also identify a rare subset of Th interneurons in mouse expressing remnant *Tac2* that has eluded capture in previous studies and is mainly localized to ventromedial striatum.

Recent studies examining the vertebrate retina at greater cell coverage have similarly identified rare mouse orthologs of populations previously considered to be primate-specific^2,32^. Future studies should address the possible functional consequences of reduced *Tac2* and increased *Th* expression in rodent striatal interneurons. In Laurasiatherians (pig and ferret), the striatal expression of *TAC3* in this ancestral initial class is conserved, but derivatives of this class also migrate to the cortex to produce an uncharacterized population. Additional studies should characterize the identity and function of the terminal cell types resulting from the reallocation of this initial class in the cortex of Laurasiatherians.

The conservation of initial classes followed by the modification of their gene expression and migratory destinations suggests that brain evolution among mammals acts on a conserved repertoire of initial classes to modify neural circuits, rather than by the creation of new cell types early in development. Emerging comparative cell atlases may yet reveal novel populations^33,34^, but it is unclear whether these populations emerge from conserved initial classes. The modification and redistribution of this initial class also raises the possibility that some initial classes may be more labile than others, representing preferred substrates for the evolutionary emergence of novel neuronal function.

## Extended data

### Supplementary Tables

**Supplementary Table 1.** Hotspot gene modules in adult and developmental datasets across species.

**Extended Data Fig. 1:**
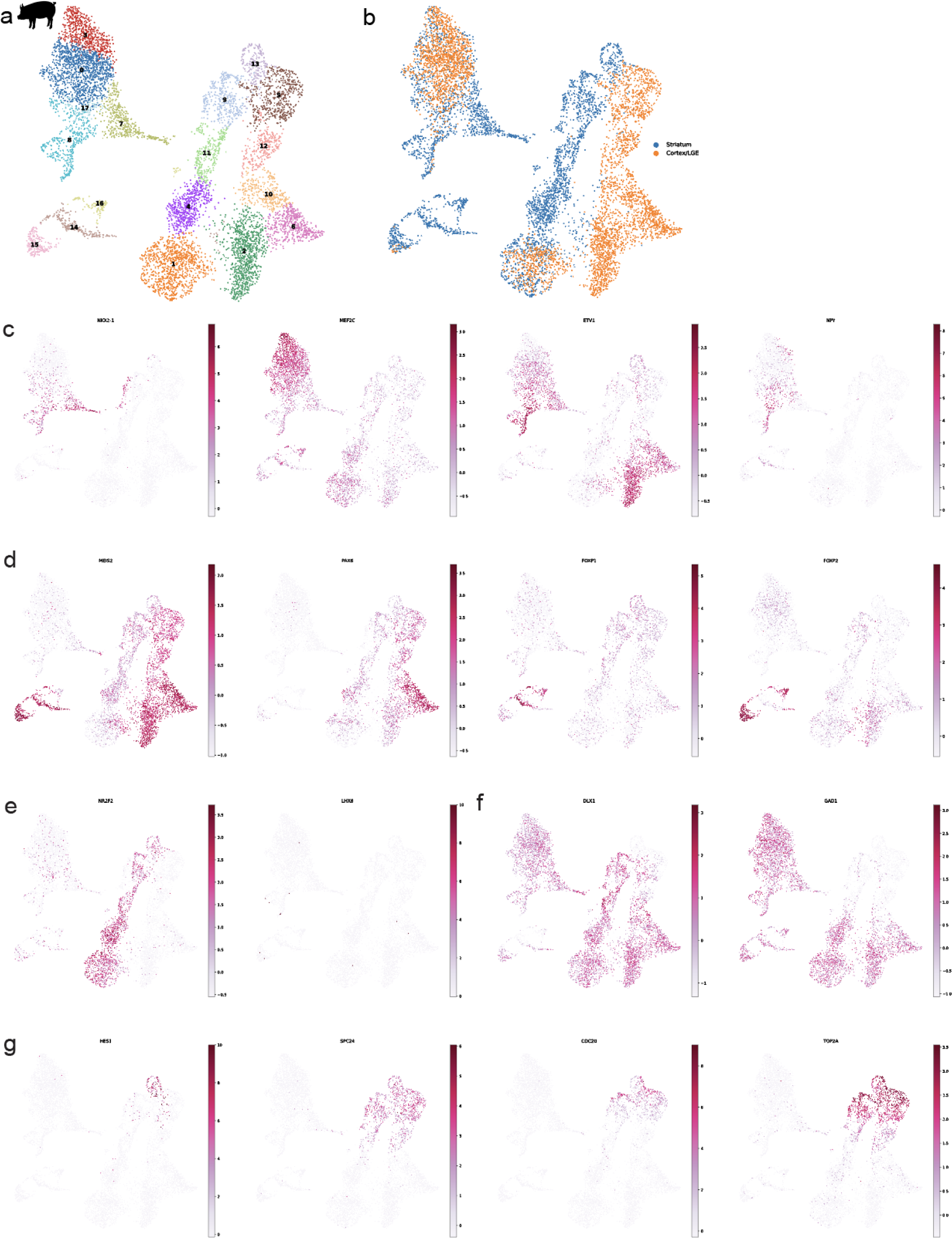
Gene expression landscape of developing pig. **a.** UMAP of Leiden clusters of E73 developing pig inhibitory neurons. **b.** UMAP colored by region of dissection. **c.** UMAP colored by conserved markers of MGE classes, scaled and normalized. NPY expression is mixed within the related MGE_LHX6/MAF class. **d.** UMAP colored by markers of LGE classes, scaled and normalized. **e.** UMAP colored by CGE and VMF markers, scaled and normalized. **f.** UMAP colored by inhibitory neuron markers, scaled and normalized. **g.** UMAP colored by markers of progenitors, scaled and normalized.

**Extended Data Fig. 2:**
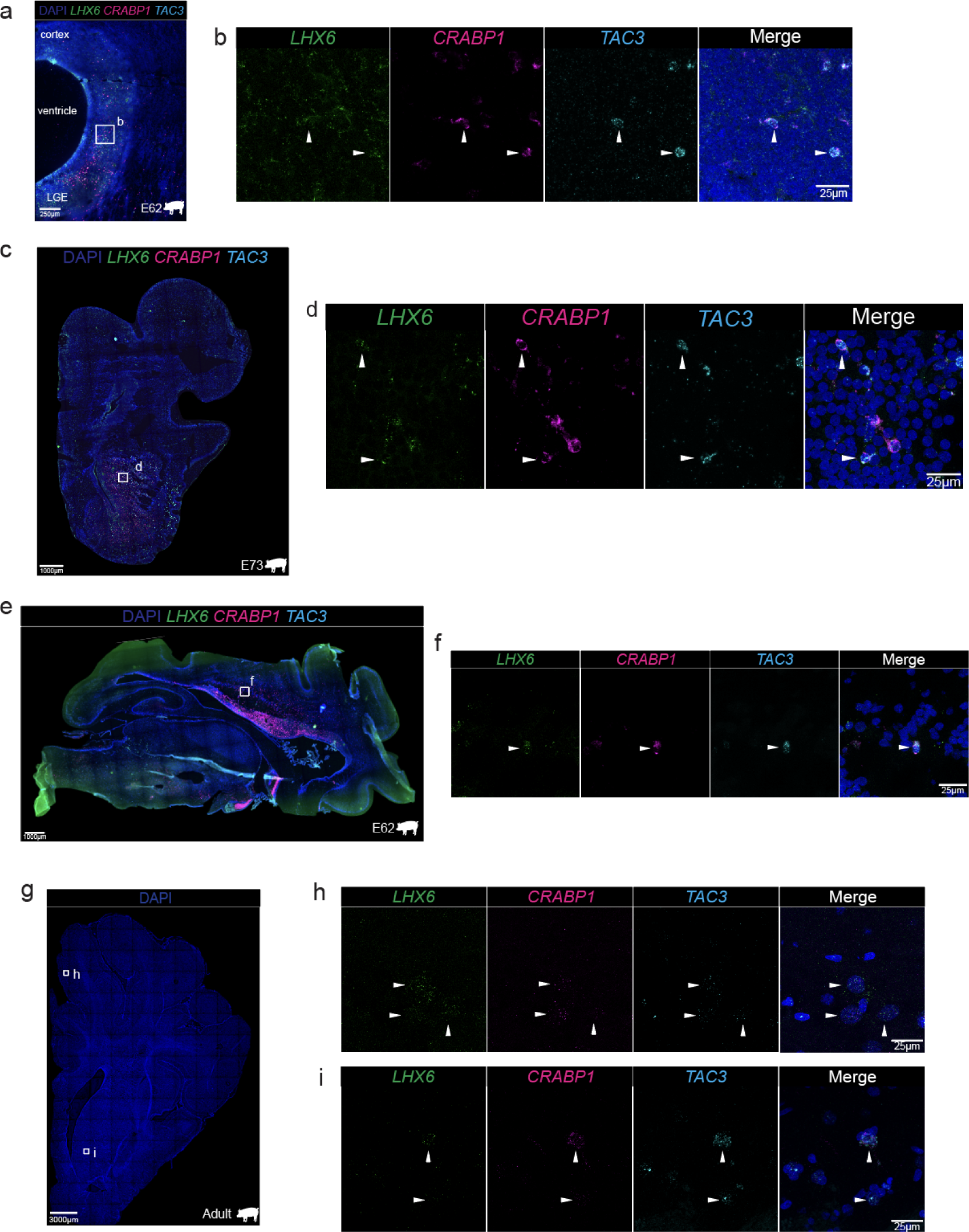
Spatial distribution of MGE_CRABP1/TAC3 class in developing and adult pig. **a.** RNAscope image of pig E62 coronal migration stream acquired at 20X with markers *LHX6, CRABP1, TAC3* with box highlighting migrating cells (b). **b.** High magnification (100X) maximum intensity projection with arrows indicating MGE_CRABP1/TAC3 class. **c.** Whole section image acquired at 10X of E73 pig coronal section RNAscope with markers *LHX6, CRABP1* and *TAC3*, with box highlighting striatum (d). **d.** High magnification (100X) maximum intensity projection of RNAscope with arrows indicating MGE_CRABP1/TAC3 initial class. **e.** Whole section image acquired at 10X of E73 pig sagittal section RNAscope with markers *LHX6, CRABP1* and *TAC3* and box highlighting cortex (f). **f.** High magnification (100X) maximum intensity projection, with arrows indicating MGE_CRABP1/TAC3 class. **g.** Whole section image acquired at 5X of the adult pig section, with boxes highlighting regions in the cortex (h), and striatum (i). **h,i.** High magnification (100X) maximum intensity projection of RNAscope with arrows indicating MGE_CRABP1/TAC3 class.

**Extended Data Fig. 3:**
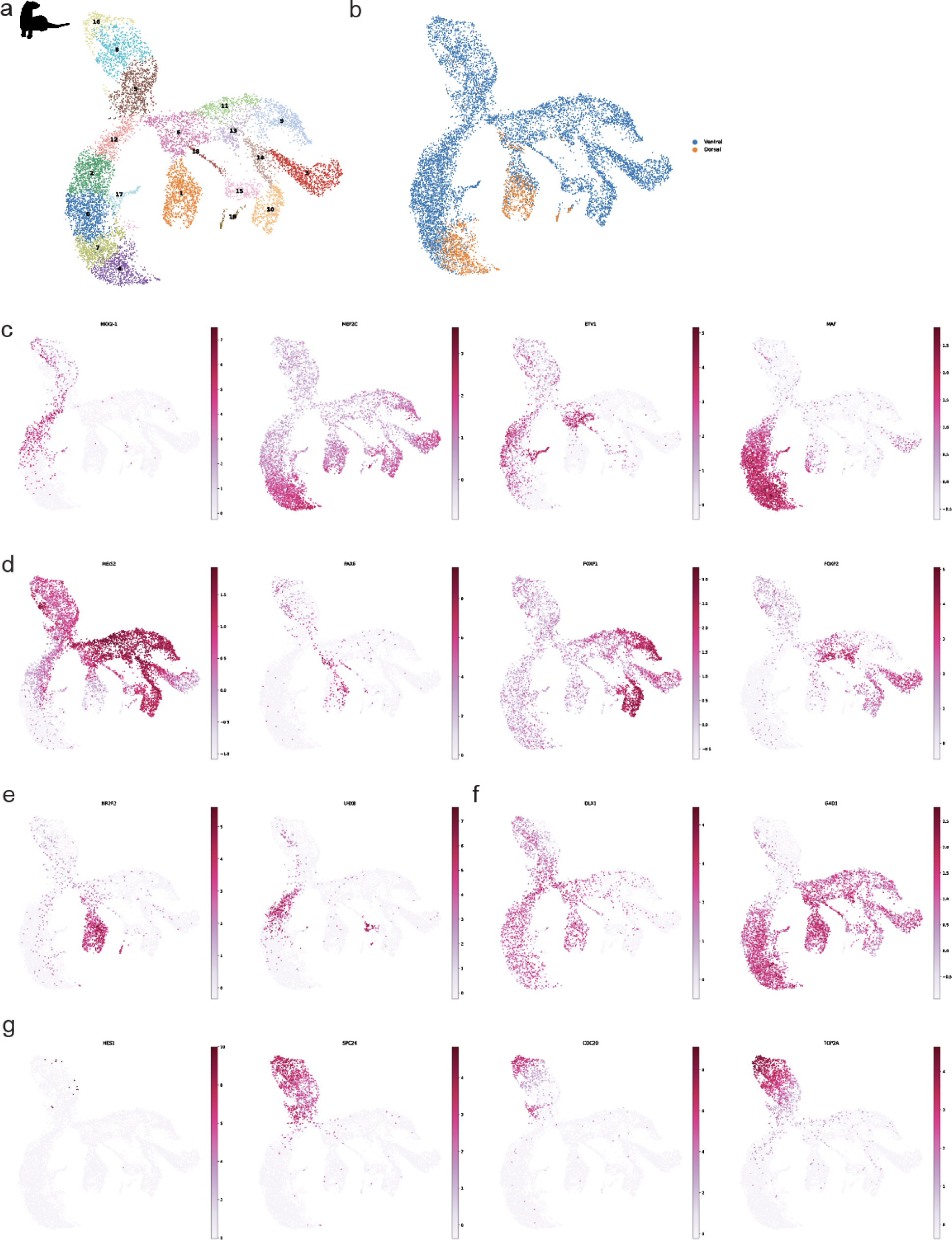
Gene expression landscape of developing ferret inhibitory neurons. **a.** UMAP of Leiden clusters from inhibitory neurons in the P1 developing ferret brain. **b.** UMAP colored by the region of dissection. **c.** UMAP colored by conserved markers of MGE classes, scaled and normalized. **d.** UMAP colored by markers of LGE classes, scaled and normalized. **e.** UMAP of CGE and VMF markers, scaled and normalized. **f.** UMAP of inhibitory neuron markers, scaled and normalized. **g.** UMAP colored by markers of progenitors, scaled and normalized.

**Extended Data Fig. 4:**
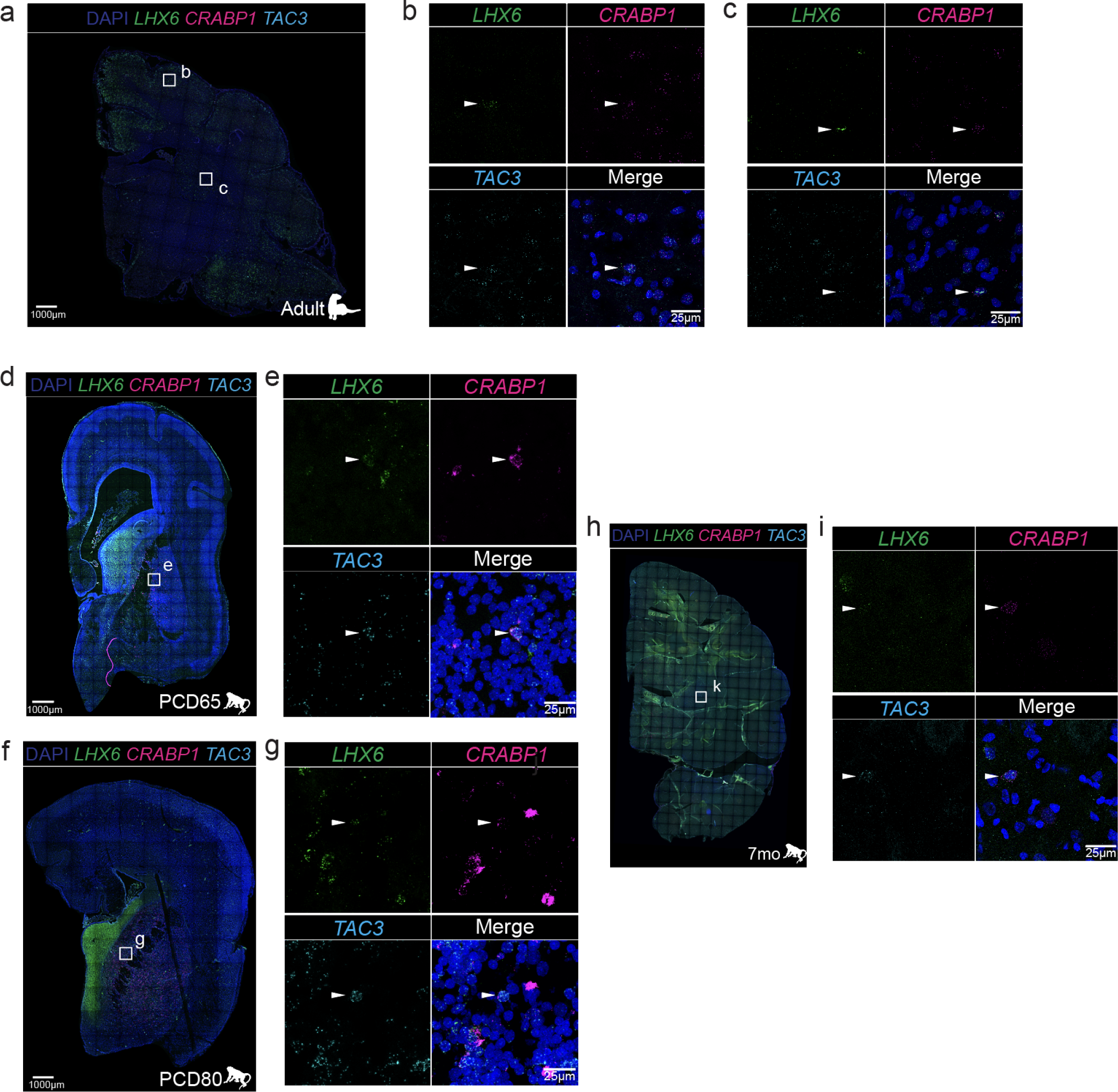
Spatial distribution of MGE_CRABP1/TAC3 class in ferret and macaque. **a.** Whole section image acquired at 20x of adult ferret RNAscope with markers *LHX6*, *CRABP1*, and *TAC3*, and boxes indicating highlighted regions in the cortex (b) and striatum (c). **b,c.** High magnification (100X) maximum intensity projection with arrowheads indicating the MGE_CRABP1/TAC3 class. **d.** Whole section image acquired at 20X of macaque PCD65 RNAscope with markers *LHX6*, *CRABP1*, and *TAC3* and box indicating the highlighted region in the striatum (e) **e.** High magnification (100X) maximum intensity projection with arrowheads indicating the MGE_CRABP1/TAC3 class. **f.** Whole section image of macaque PCD80 RNAscope with markers *LHX6*, *CRABP1*, and *TAC3*, and box indicating the highlighted region in the striatum (g). **g.** High magnification (100X) maximum intensity projection with arrowheads indicating the MGE_CRABP1/TAC3 class. **h.** Whole section image acquired at 10X of 7 months old macaque RNAscope with markers *LHX6*, *CRABP1*, and *TAC3*, with boxes indicating the highlighted region in the striatum (i). **i.** High magnification (100X) maximum intensity projection with arrowheads indicating the MGE_CRABP1/TAC3 class.

**Extended Data Fig. 5:**
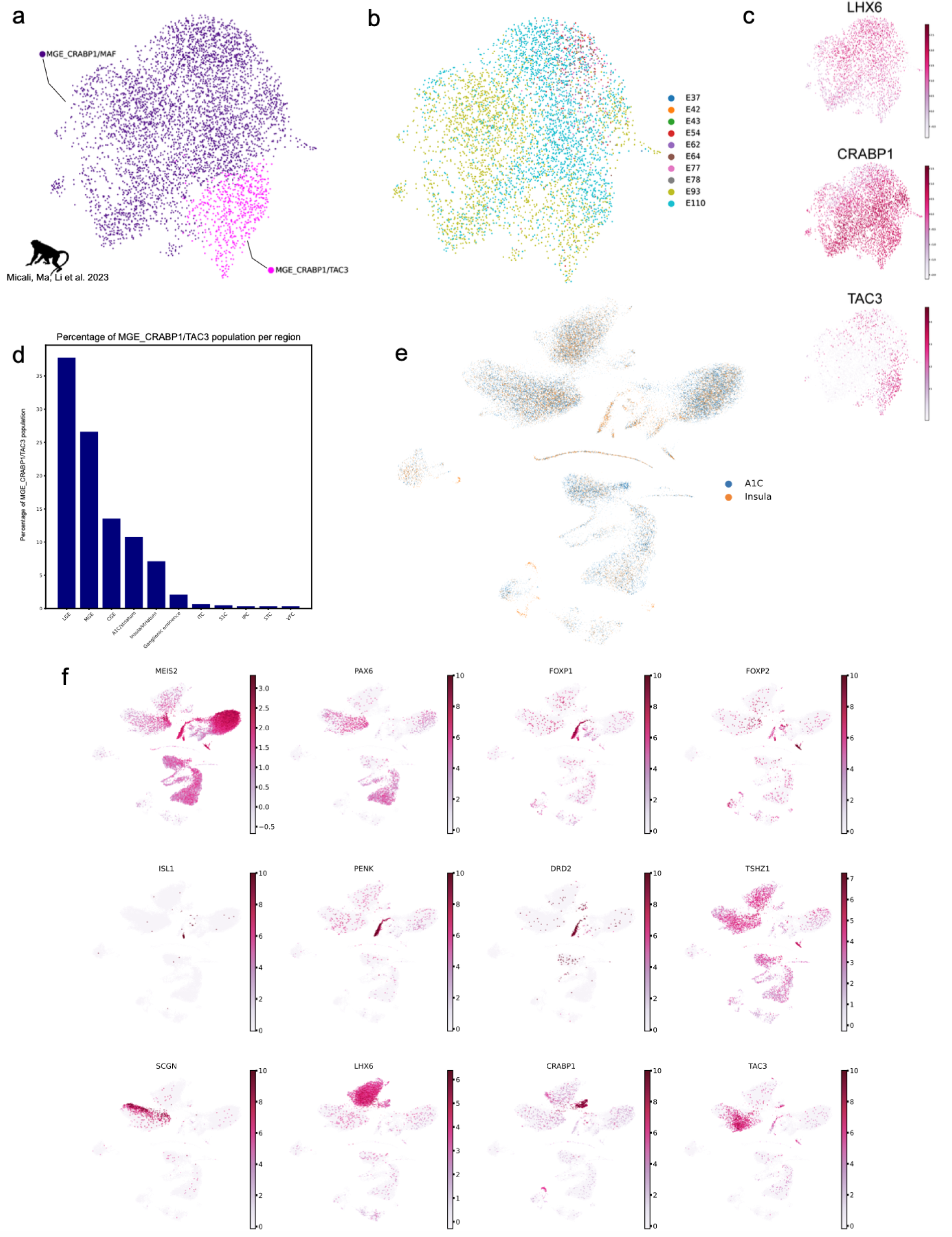
Analysis of published developmental macaque data. **a.** UMAP colored by MGE_CRABP1 initial class. **b.** UMAP colored by age of sample. **c.** UMAP colored by markers of MGE_CRABP1/TAC3 class. **d.** Bar plot representing the percentage of MGE_CRABP1/TAC3 population isolated from specified dissection regions, with A1C and Insula dissections renamed to include striatum. **e.** A1C and Insula samples isolated, normalized, scaled and clustered, colored by region of dissection as originally stated. f. Gene expression of medium spiny neurons and MGE_CRABP1/TAC3 initial class in A1C and insula dissection regions, indicating contaminating striatum.

**Extended Data Fig. 6:**
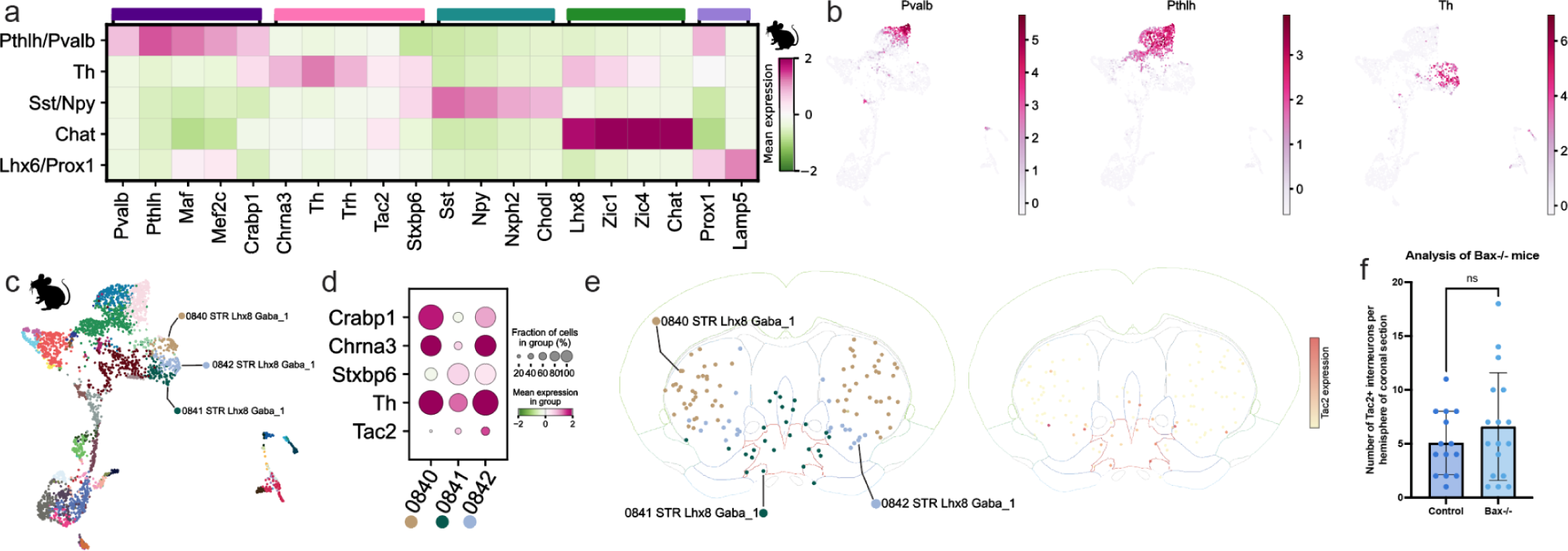
Gene expression, spatial distribution and Bax-/- analysis of adult mouse *Tac2*-expressing population. **a.** Heatmap of selected markers of inhibitory neuron terminal classes in adult mouse, scaled and normalized. **b.** UMAP of Pthlh/Pvalb and Th markers in adult mouse, scaled and normalized. **c.** UMAP colored by Allen Brain Cell Atlas (ABC Atlas) cluster annotation. **d.** Dotplot of *Tac2+* Th interneuron marker gene expression in highlighted ABC Atlas clusters, normalized and scaled. **e.** Distribution of highlighted ABC clusters in the adult mouse brain and *Tac2* expression found within those populations. Image credit: Allen Brain Cell Atlas (RRID:SCR_024440) https://portal.brain-map.org/atlases-and-data/bkp/abc-atlas. **f.** Number of *Lhx6*, *Crabp1*, *Tac2* expressing cells per hemisphere in 20µm section of P15 Bax^-/-^ (n=17) and control (Bax^-/+^)(n=14) sample, ‘ns’ indicates difference is not significant (p=0.3037, t=1.049).

**Extended Data Fig. 7:**
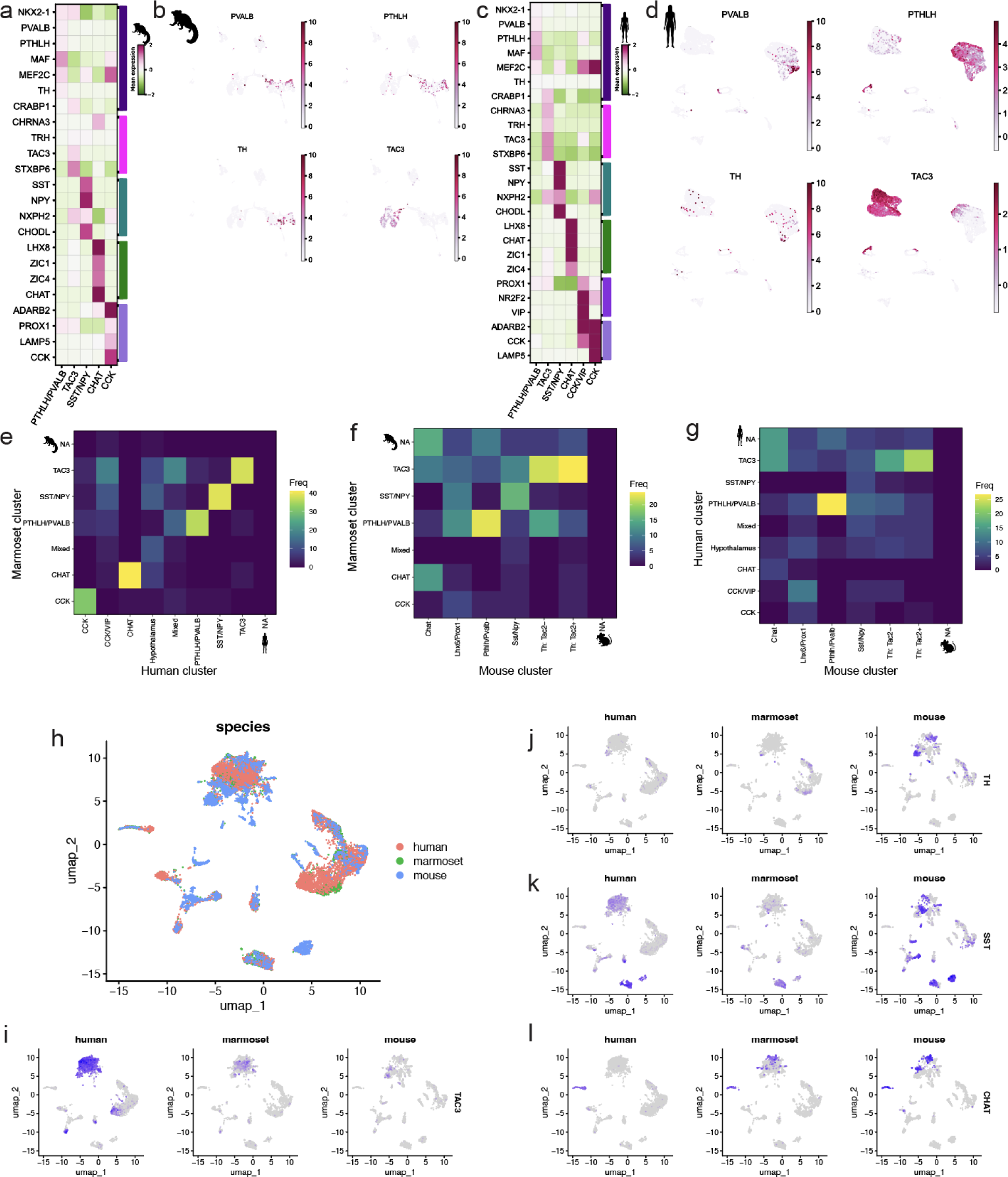
Gene expression landscape of adult human and marmoset striatal interneurons and LIGER integration of adult single nucleus RNA sequencing data. **a.** Heatmap of selected markers of striatal inhibitory neuron terminal classes in adult marmoset, scaled and normalized. **b.** UMAP colored by markers of inhibitory neuron terminal classes in adult marmoset, scaled and normalized. **c.** Heatmap of selected markers of inhibitory neuron terminal classes in adult human, scaled and normalized. **d.** UMAP colored by markers of inhibitory neuron terminal classes in adult human, scaled and normalized. **e-g.** LIGER co-clustering frequencies across 42 parameter combinations using 500 variable features (see methods). ‘NA’ denotes lack of co-clustering across species. **h.** Representative Liger integration of human, marmoset, and mouse data, colored by species (500 features, k=30, lambda =8).**i-l.** Liger integration UMAP colored by selected markers of inhibitory interneurons in each species.

**Extended Data Fig. 8:**
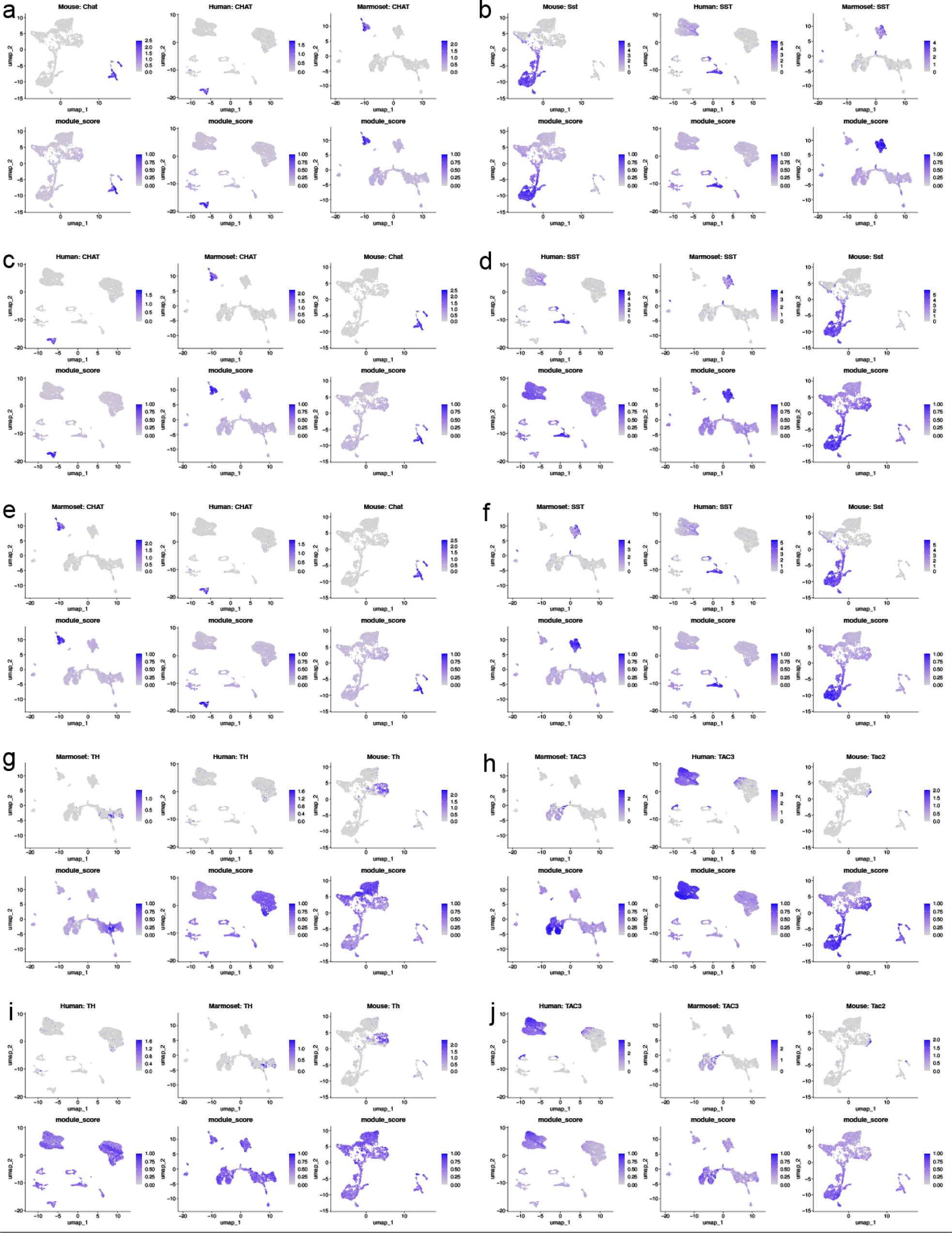
Hotspot gene modules across adult datasets a,b. Mouse *Chat* and *Sst* Hotspot modules projected onto mouse, human and marmoset. **c,d.** Human *CHAT* and *SST* Hotspot modules projected onto human, marmoset and mouse. **e,f.** Marmoset *CHAT* and *SST* Hotspot modules projected onto marmoset, human and mouse. **g,h.** Marmoset *TH* and *TAC3* Hotspot modules projected onto marmoset, human and mouse. **i,j.** Human *TH* and *TAC3* Hotspot modules projected onto human, marmoset, and mouse.

**Extended Data Fig. 9:**
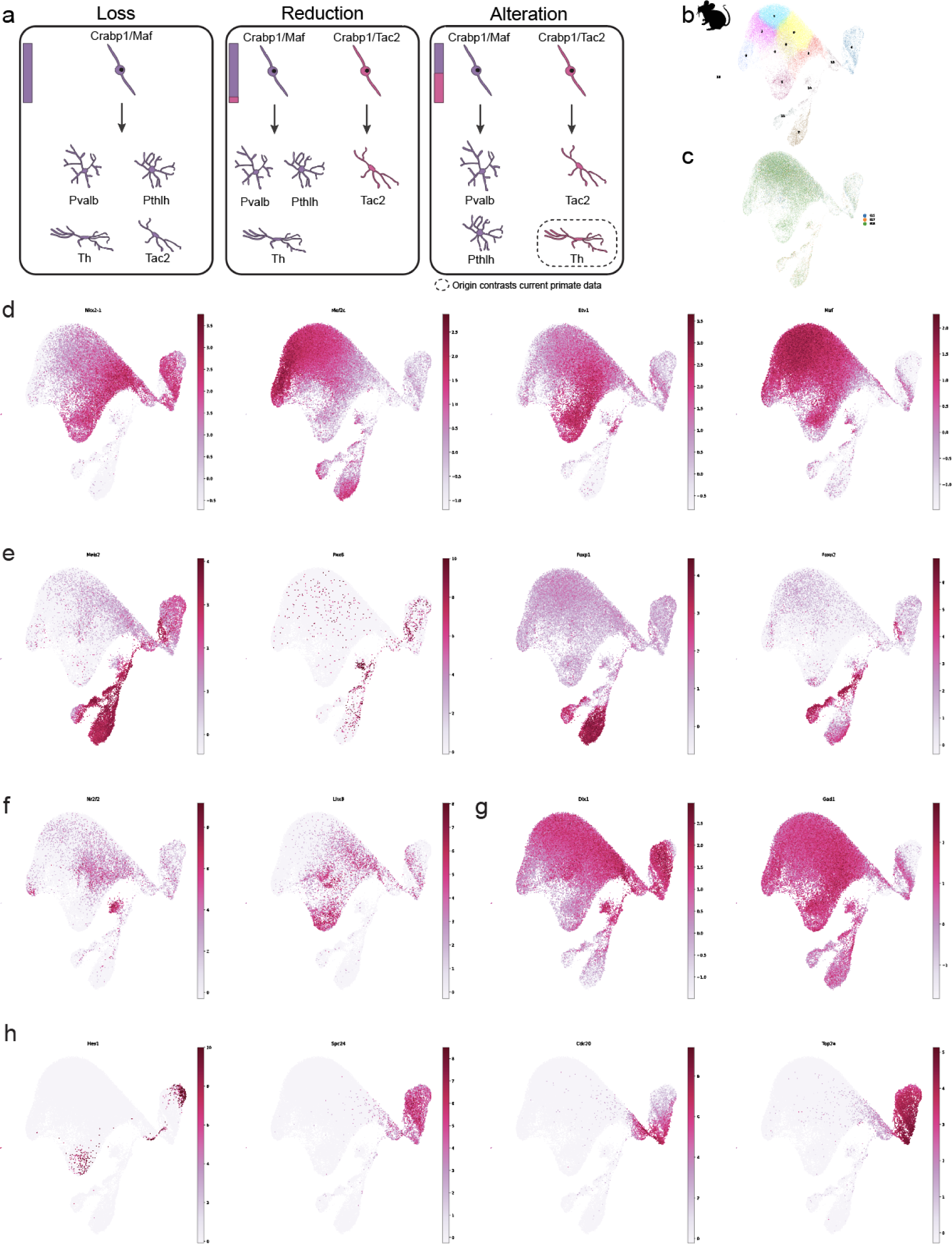
Gene expression landscape of developing mouse. **a.** Schematic of developmental scenarios that could lead to small *Tac2* expressing population and differences in *Th* expression between primate and mouse striatal interneurons. Bars indicate relative abundance of initial classes. **b.** UMAP colored by Leiden clusters of inhibitory neurons and their progenitors in the developing mouse MGE/LGE/striatum. **c.** UMAP colored by developmental timepoint. **d.** UMAP colored by conserved markers of MGE classes, scaled and normalized. **e.** UMAP colored by markers of LGE classes, scaled and normalized. **f.** UMAP of CGE and VMF markers, scaled and normalized. **g.** UMAP of inhibitory neuron markers, scaled and normalized. **h.** UMAP colored by markers of progenitors, scaled and normalized.

**Extended Data Fig. 10:**
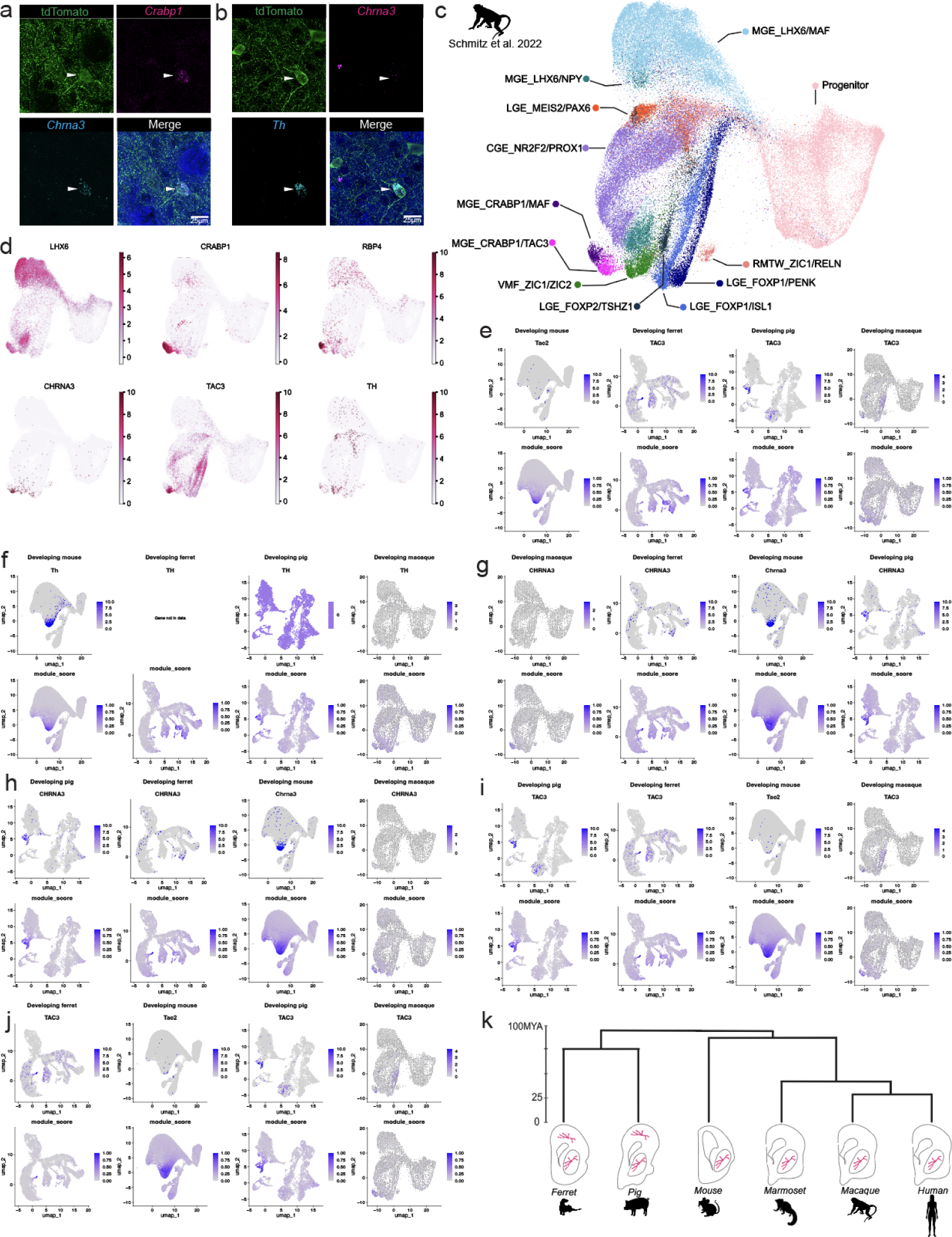
Comparison of inhibitory neuron initial classes across mouse, macaque, pig, and ferret. **a.** High magnification (100X) maximum intensity projection of adult Nkx2.1-Cre;Ai14 mouse brain tdTomato reporter expression and *Crabp1*, *Chrna3* RNAscope with arrowhead indicating the MGE_CRABP1/CHRNA3 initial class. **b.** High magnification (100X) maximum intensity projection of adult Nkx2.1-Cre;Ai14 mouse brain tdTomato reporter expression and *Chrna3*, *Th* RNAscope with arrowhead indicating Th interneuron. **c.** UMAP colored by inhibitory initial classes in developing macaque scRNAseq data from Schmitz et al. 2022. **d.** UMAP colored by markers of *LHX6/CRABP1* initial classes, scaled and normalized. **e,f.** Developing mouse *Tac2* and *Th* Hotspot modules project to MGE_CRABP1/TAC3 in ferret, pig and macaque. **g.** Developing macaque *CHRNA3* Hotspot module lacked the resolution to differentiate between *CRABP1*-expressing classes in macaque. **h.** Developing pig *CHRNA3* Hotspot module. **i.** Developing pig *TAC3* Hotspot module. **j.** Developing ferret *TAC3* Hotspot module lacked the resolution to differentiate between *CRABP1*-expressing classes in macaque and mouse. **k.** Schematic of phylogenetic and spatial distribution of MGE_CRABP1/CHRNA3 initial class.

## Methods

### Mouse experimental model and subject details

All animal procedures conducted in this study followed experimental protocols approved by the Institutional Animal Care and Use Committee of the University of California, San Francisco (UCSF). The Nkx2.1-Cre;Ai14 mouse line is the result of crossing the C57BL/6J-Tg(Nkx2.1-cre)2Sand/J (“Nkx2.1-Cre”; Jackson Laboratory stock no. 008661) and B6.Cg-*Gt(ROSA)26Sor^tm14(CAG-tdTomato)Hze^*/J (“Ai14”; Jackson Laboratory stock no. 007914) strains. Mouse housing and husbandry were performed in accordance with the standards of the Laboratory Animal Resource Center (LARC) at UCSF. Mice were group housed in a 12 h light/dark cycle, with access to food and water *ad libitum*.

Nkx2.1Cre;Ai14 mice were crossed, and the date of a positive vaginal plug was considered as E0. Pregnant dams were sacrificed, and the embryonic brains from their litters were extracted and assessed for tdTomato fluorescence, at the following developmental stages: E15 (n=16 embryos from two litters), E17 (n=7 embryos from one litter) and E18 (n=5 embryos from two litters). The LGE/striatum and the mantle zone of the MGE were manually dissected and collected for further processing. The entire dissection process was carried out on ice, using Hibernate-E culture medium. The sex of the embryos was not determined, and thus the results reported are assumed to include animals of both sexes.

Dissected regions were incubated with a prewarmed solution of papain (Worthington Biochemical Corporation) prepared according to the manufacturer’s instructions. After 30–60 min of incubation, samples were gently triturated with wide orifice pipette tips. Once the samples were dissociated to a single cell suspension, DMEM + 0.1% BSA (Sigma) was added to quench papain and cells were pelleted at 300g for 5 minutes at 4°C. Cells were resuspended in PBS supplemented with 0.1% BSA and sorted for TdTomato expression on a BD FACSAria Fusion.

### Pig and ferret samples

Ferret samples from P1 and adult (33 months of age) were a gift from the University of Iowa National Ferret Resource and Research Center. Pig samples from E62, E73, and adult (1 year and 2 months of age) were collected at the Swine Teaching and Research Center at the University of California, Davis. All animal procedures were performed in accordance with the standards of the Animal Welfare Act, carried out under the Association of Assessment and Accreditation of Laboratory Animal Care International (AAALAC) approved conditions and followed experimental protocols approved by Institutional Animal Care and Use Committee (IACUC) at the University of California, Davis.

For the P1 ferret sample, dissection was performed in Hibernate-E (Gibco A1247601) under a stereo dissection microscope (Olympus SZ61) on ice. The brain was bisected and cut coronally to reveal MGE, LGE, striatum, and cortex. Cortex was processed independently from ventral regions. For the E73 pig sample, the brain was embedded in low-melting point agarose and vibratome sectioned at 300 uM in ACSF (125 mM NaCl, 2.5 mM KCl, 1 mM MgCl_2_, 1 mM CaCl_2_, and 1.25 mM NaH_2_PO_4_). Following sectioning, tissue sections were microdissected under a stereo dissection microscope in Hibernate-E on ice to capture striatal and cortical populations. For single-cell dissociation of pig and ferret samples, dissected regions were cut into small pieces and incubated with a prewarmed solution of papain. After 30–60 min of incubation, samples were gently triturated with wide orifice pipette tips. Once the samples were dissociated to a single cell suspension, DMEM + 0.1% BSA was added to quench papain and cells were pelleted at 300g for 5 minutes at 4°C. Finally, cells were resuspended in PBS supplemented with 0.04% BSA.

### Single cell sequencing of developing mouse, pig, and ferret

Single-cell RNA sequencing was completed using the 10x Genomics Chromium X controller and version 3.1 high-throughput RNA capture kits. Samples were loaded at approximately 30,000-100,000 cells per well and library preparation was completed following manufacturer’s instructions. Libraries were sequenced on Illumina HiSeq and NovaSeq X platforms. Sequencing was performed at the UCSF CAT, supported by UCSF PBBR, RRP IMIA, and NIH 1S10OD028511-01 grants.

### Alignments and gene models

Illumina BCL files were converted to Fastq files using bcl2fastq2. Genes were quantified using CellRanger v7.0.1 count function using the MusPutFur1.0 genome for the ferret sample, SusScrofa11.1 for the pig sample, and mm10-2020A for the mouse. To improve recovery of missing transcripts, we utilized the ReferenceEnhancer^35^ package to optimize the SusScrofa and MusPutFur genomes by incorporating intergenic reads and manually annotating missing TAC3 interneuron marker genes.

### Quality control

CellRanger output was input into Cellbender v0.2.1^36^ to remove ambient RNA and Cellbender filtered output matrix was used for downstream processing. For pig data processing, droplets with fewer than 1750 genes and greater than 6000 were filtered from the dataset and droplets with greater than 10% mitochondrial and ribosomal reads were also removed from the dataset. Doublets were detected and removed using Scrublet (version 0.2.3, threshold set to 0.25)^37^. For ferret data processing, droplets with fewer than 1000 or greater than 6000 genes and greater than 6% ribosomal were removed from the dataset. Doublets were removed using Scrublet. For mouse data processing, droplets with fewer than 2250 genes and greater than 2% ribosomal and mitochondrial reads were removed. Doublets were removed using Scrublet.

### Clustering and assignment of cell types

Analysis was based upon the Scanpy package and tutorial^38^. Counts were normalized and log transformed. The data was then scaled for each gene and principal-component analysis was performed. Batch correction was applied to the pig and mouse data using Harmony integration^39^. Leiden clustering was applied with resolution set to 1. Inhibitory neurons were then isolated as previously described^6^. Data was reset to raw, and normalization, log transformation, scaling, principal-component analysis and Leiden clustering at resolution of 1 or 0.5 (mouse) were repeated on the isolated inhibitory neurons. Individual Leiden clusters were subdivided by increasing resolution for individual clusters or merged to initial classes. Initial classes and nomenclature were assigned based on markers in previous work^6^.

### Analysis of published developing macaque dataset

Data was taken from Micali, Ma, Li, et al^24^. We first isolated the LHX6_CRABP1 population, normalized, log transformed, scaled, batch corrected, and performed Leiden clustering. Clusters were labeled based on gene expression of known markers. Due to the presence of MGE_CRABP1 populations in the A1C and Insula dissection, we next isolated cells from A1C and Insula dissections. We again normalized, log transformed, scaled, batch corrected, and performed Leiden clustering. We examined marker gene expression and renamed dissections A1C and Insula to include striatum due to the presence of medium spiny neurons in these dissections.

### Analysis of published adult marmoset, human, and mouse datasets

Adult mouse striatal interneurons cells were curated from the Yao et al. 2023 Allen Brain Cell (ABC) atlas class “08 CNU-MGE GABA” (n = 18849 cells), which we subset further to include only striatal cells (anatomical_division_label = “STR”), comprised of 4785 cells derived from a single ABC dataset (“WMB-10Xv3-STR”). This dataset initially contained cells from ABC subclasses “054 STR Prox1 Lhx6 Gaba”, “055 STR Lhx8 Gaba”, “056 Sst Chodl Gaba”, “057 NDB-SI-MA-STRv Lhx8 Gaba”, and “058 PAL-STR Gaba-Chol”, but upon further examination, we found that majority of cells from ABC subclass 057 originate from outside the striatum. Exclusion of subclass 057 resulted in a final dataset of 3586 cells.

Adult human striatal interneurons were curated from the neuron dataset of the Siletti et al. 2023 atlas by first selecting for the ROI terms “Basal Nuclei (BN) - Body of the Caudate – CaB”, “Basal Nuclei (BN) -Putamen – Pu”, and “Basal Nuclei (BN) - Nucleus Accumbens – NAC”, totaling 88044 cells. These cells were further subset to include only cells from supercluster terms “CGE interneuron” (n = 711), “MGE interneuron” (n = 238), or “Splatter” (n = 6716). The remaining 7665 cells were re-embedded (UMAP from 30 PCs) using 2000 variable features (Seurat “vst”) and re-clustered using Louvain (at arbitrary resolution 0.8) into 22 clusters. These clusters were used to further exclude putative MSNs based on MEIS2 expression, producing a final dataset of 6617 striatal interneurons.

Adult marmoset striatal interneurons were curated from the Krienen et al. 2023 Marmoset Census dataset of 6249 striatal GABAergic neurons (“striatum.GAD”). Again we excluded MEIS2-expressing cells (in this case using marmoset census cluster “02”). Using the same method as adult human data, the remaining 3930 cells were re-embedded and clustered into 22 clusters, from which we identified 6 small clusters (totaling 551 cells) that appeared to be contamination from excitatory (SLC17A6- and SLC17A-expressing) neurons (possibly from doublets), which we excluded, leaving 3379 cells in the final dataset.

Following cell selection in each species, the data was reprocessed using a typical Seurat (v5.0.3) pipeline: NormalizeData (natural log of [1 + reads per 10-thousand cells]), FindVariableFeatures (2000 features using Seurat “vst”), ScaleData/PCA, FindNeighbors (using 30 PCs), FindClusters (Louvain resolution 0.8), and RunUMAP (using 30 PCs). In all cases we found our re-embeddings were consistent/coherent with the previously published cell groupings/labels.

### Cross-species integration of adult datasets

For cross-species integrations, we utilized LIGER^29^ (R package rliger v.1.0.1) to perform integration based on iNMF (inverse non-negative matrix factorization). We chose LIGER iNMF after verifying that conventional batch-correction integration approaches (Seurat CCA, JPCA, RPCA, Harmony, and scVI) largely fail to group cells across species (data not shown).

For LIGER integrations, we identified a set of gene orthologs across human, mouse, and marmoset using ENSEMBL v110. Of genes with annotated 3-way orthology, we subset to only genes that are also found within the 3 original datasets, removing one additional gene for which 1 mouse gene was assigned 2 primate orthologs, and adding back primate TAC3/mouse Tac2 (as this orthology was not included in ENSEMBL v110), which left a final matched set of 9875 genes.

Using these genes and our curated adult datasets, we generally followed the recommendations of the LIGER authors^40^.After merging the 3 datasets and re-normalizing on per-cell basis, we performed integrations using 500, 1000, and 2000 variable features (Seurat “vst”), in each case re-scaling the data (splitting by species but not re-centering i.e., using Seurat ScaleData with split.by=”species” and do.center=FALSE to divide each gene by its root mean square). For each set of variable features, we used 42 combinations (for 126 total integrations) of the LIGER parameters *k* (which correlates with dimensionality) and *lambda* (a penalty for species-specific clustering). Following an elbow estimation from the LIGER function ‘suggestK’, we explored *k* values in 20, 25, 30, 35, 40, and 45. Given the uncertainty inherent to determining the species-specificity of any given cell-type, we explored a wide range of *lambda* values: 0.5, 1, 2, 4, 8, 16, and 32. In each case, we ran LIGER RunOptimizeALS and RunQuantileNorm (each splitting by species), Seurat FindNeighbors using the first 20 iNMF dimensions, Seurat RunUMAP using *k* iNMF dimensions, and we found integrated clusters using Louvain at resolution 0.5.

To quantify co-clustering within an integration for some set of subject cells e.g., the 51 mouse Tac2-expressing cells, we identified the integrated cluster into which the subject cells most frequently reside. Then, for some query annotation e.g., human cell clusters, we identify the query annotation most frequently observed within that integrated cluster. We then aggregate this across the 126 integrations. In each reported case, we saw no obvious relationship between either parameter *k* or *lambda* and the most frequent co-clustering result, but 500 features generally produced the most consistent co-clustering results.

### Gene module discovery and projection using Hotspot

Hotspot (hotspotsc v1.1.1; https://github.com/YosefLab/Hotspot) was used for unsupervised partitioning of genes into modules following the authors’ recommendations^30^. For each dataset, only genes present with nonzero counts were used, and Hotspot was run using model “danb” on the PCA embedding. Genes were selected from Hotspot autocorrelations with FDR < 0.05 and partitioned using agglomerative clustering, as previously demonstrated by the authors using the default ‘min_gene_threshold=30’.

To address computational constraints on hotspot module discovery in the larger developing macaque (109111 cells) and developing mouse (71023 cells) datasets, we subset 50% of the cells. To make the subsets representative, we sampled within high-resolution clusters: for macaque, we used the 124 ‘hires_leiden’ clusters already present in the data; for mouse, we re-clustered (leiden at resolution=10) to obtain 157 distinct clusters used only for this purpose. All other analyses (including module scores/projections) use the full datasets.

Gene module scores were calculated using PC1 following PCA using only the genes within the module. For cross-species module projections, we found the unsupervised gene module most correlated (Spearman) with a given marker gene of interest. We then found the gene orthologs in other species (subsetting genes if necessary) and calculated module scores.

### Immunohistochemistry tissue processing

P1 and adult ferret, pig, Bax and developing mouse samples were fixed in 4% paraformaldehyde in PBS overnight at 4 °C with constant agitation. The paraformaldehyde was then replaced with fresh PBS (pH 7.4) and samples were cryopreserved by incubation for 24 h in 10% sucrose diluted in PBS (pH 7.4), followed by a 24 h incubation in 20% sucrose, and finally a 24 h incubation in 30% sucrose before being embedded in OCT (Tissue-Tek, VWR). Tissue was then frozen at –80 °C, cryosectioned at 20 μm and stored at -80°C until used. Slides from P22 ferret were a gift from A. Kriegstein (University of California San Francisco).

Adult Nkx2.1-Cre;Ai14 mice were transcardially perfused with PBS followed by 4% PFA in PBS; their brains were dissected out and postfixed overnight at 4°C in 4% PFA in PBS with constant agitation. 50-µm sections were obtained on a vibratome, and preserved in freezing buffer (30% ethylene glycol, 28% sucrose in 0.1 M sodium phosphate buffer) at -20°C until used.

Rhesus macaque sections from previous studies were provided by the Primate Center at the University of California, Davis and prepared for histology as stated above. All animal procedures conformed to the requirements of the Animal Welfare Act, and protocols were approved before implementation by the Institutional Animal Care and Use Committee at the University of California, Davis.

RNAscope was performed following manufacturer’s instructions for the Advanced Cell Diagnostics RNAscope Multiplex Fluorescent Reagent Kit V2 Assay (ACD, 323120).For immunostaining of Nkx2.1-cre;Ai14 line, rabbit anti-RFP (Rockland 600-401-379) was diluted 1:1000 in blocking buffer (PBS + 5% BSA + 0.3% Triton X-100) and incubated for 1 hour at 37 °C. Alexa dye-conjugated goat secondary 594 antibody was diluted 1:500 in blocking buffer and incubated for 30 minutes at 37°C.

After RNA *in situ* hybridization, TrueBlack lipofuscin autofluorescence quencher (Biotium, 23007) was applied to adult ferret, adult pig and 7 month old macaque samples according to the manufacturer’s directions. Slides were mounted using Prolong Gold Antifade plus Dapi (Invitrogen, P36931).

### Analysis of Bax-/- mice

P15 male CD1 Bax^tm1Sjk^ (MGI: 1857429) were a gift from D. Laird (University of California San Francisco). Heterozygous litter matched samples were used as control. RNAscope was performed on 20 μm sections using markers *Lhx6, Crabp1*, and *Tac2*. Whole tissue section images were acquired using a 20x objective. Scans were imported into Imaris v10.1.0 (Bitplane) and triple positive cells were manually marked using the Spots tool then manually counted per hemisphere. Quantification was compared statistically using T-test with Welch’s Correction.

### Imaging

Entire tissue sections were imaged single plane in widefield mode on a Leica DMI8 inverted microscope (Leica Microsystems) connected to a Flash4 V3 camera (Hamamatsu) camera, with either a 5x (0.12NA), 10X (0.4NA) or 20X (0.8) Plan Apochromat objective. Stitching was performed automatically in LASX. Regions of interest were imaged at higher resolution using a laser scanning confocal microscope (Stellaris, Leica) 100X oil (1.4NA) Plan Apochromat objective with a pixel size of 116.25µm x 116.25µm. Z-stacks of the entire volume of cells were acquired using optimal sections. Images from all samples were acquired under the same imaging settings with minimal adjustment of illumination intensity.

### Image Processing

Maximum intensity projections were produced from volumes acquired on the confocal microscope, then brightness and contrast were manually adjusted in Fiji for publication. Image processing of scans was performed using Imaris v10.1.0 (Bitplane). Files were converted using the Imaris v10 Converter and imported into Surpass mode for interaction. Images were saved as 3500 dpi PNG files for publication.

## Data availability

The sequencing data will be made available upon publication. Scripts and annotation files for the study have been deposited on github at https://github.com/mdeber/Corrigan2024.

## Acknowledgements

We thank M. Song, A. Kriegstein, J. Zussman, D. Laird, D. Glawe, M. Starkenburg, E. Maga, P. Ji, S. Sopocy, C. Sehnert, A. Taranatal, and A. Fox for samples for genomic and histological analysis. We thank M. Mui, C. Mrejen and E. Gaylord for assistance in sample preparation, imaging, and image analysis, respectively. Imaging was performed at Innovation Core at the Weill Institute for Neurosciences. This material is based upon work supported by the National Science Foundation under grant number 2034836 (E.K.C.). This work was supported by the following funding sources: National Institutes of Health DP2MH122400, U01MH114825, UM1MH130991, and Schmidt Futures Foundation. AAP is a New York Stem Cell Foundation Robertson Investigator.

## Author contributions

E.K.C., M.F.P., F.M.K. and A.A.P. designed the methodology. E.K.C., A.P., M.T.G. performed the experiments. E.K.C. conducted microscopic analysis. E.K.C., A.P., M.T.G. performed the single-cell RNA sequencing processing. E.K.C performed single-cell RNA sequencing analysis of developmental datasets. M.D. performed single-cell RNA sequencing analysis of published adult datasets. E.K.C. and A.A.P. wrote the manuscript with input from all authors. E.K.C., C.C.H., M.F.P., C.C.K. and A.A.P. acquired the funding. E.K.C., M.F.P., F.M.K., and A.A.P. conceptualized the study. M.T.S., C.C.H., M.F.P., F.M.K., A.A.P. supervised the project

